# Comparative reproductive biology of deep-sea ophiuroids inhabiting polymetallic-nodule fields in the Clarion-Clipperton Fracture Zone

**DOI:** 10.1101/2021.02.06.428832

**Authors:** Sven R Laming, Magdalini Christodoulou, Pedro Martinez Arbizu, Ana Hilário

**Affiliations:** Centre for Environmental and Marine Studies (CESAM) & Department of Biology, University of Aveiro, 3810-193 Aveiro, Portugal; German Centre for Marine Biodiversity Research (DZMB), Senckenberg am Meer, 26382 Wilhelmshaven, Germany

**Keywords:** maturation, lecithotrophy, gonochoric, deep-sea mining, ecology, brittle stars, Ophiuroidea, Echinodermata

## Abstract

Deep-sea mining in the Pacific Clarion-Clipperton Fracture Zone (CCZ), a low-energy sedimentary habitat with polymetallic nodules, is expected to have considerable and long-lasting environmental impact. The CCZ hosts extraordinarily high species diversity across representatives from all Domains of Life. Data on species biology and ecology remain scarce, however. The current study describes the reproductive biology of *Ophiosphalma glabrum* (Lütken & Mortensen, 1899) (Ophiosphalmidae) and *Ophiacantha cosmica* (Lyman, 1878) (Ophiacanthidae), two ophiuroids frequently found in the CCZ. Specimens collected in Spring 2015 and 2019 in four contract areas were examined morphologically and histologically. Size-class frequencies (disc diameter and oocytes feret diameters), sex ratios, gametogenic status, putative reproductive mode and a simple proxy for fecundity are presented. Habitat use differs in each. While *Ophiosphalma glabrum* is epibenthic, occurring as single individuals, *Ophiacantha cosmica* often form size-stratified groups living on stalked sponges, suggesting gregarious settlement or retention of offspring (though no brooding individuals were found). Further molecular analyses are needed to establish whether *O. cosmica* groups are familial. In *Ophiosphalma glabrum*, for which sample sizes were larger, sex ratios approximated a 1:1 ratio with no size-structuring. In both species, individuals were at various stages of gametogenic maturity but no ripe females were identified. Based on this, *O. glabrum* is most probably gonochoric. Reproductive mode remains inconclusive for *Ophiacantha cosmica*. Both species are presumptively lecithotrophic, with vitellogenic-oocyte feret diameters exceeding 250 µm. Oocyte feret diameters at times exceeded 400 µm in *Ophiosphalma glabrum*, indicating substantial yolk reserves. Estimates of instantaneous fecundity (vitellogenic specimens of *O. glabrum* only) were confounded by interindividual variability in gonad characteristics. The well-furnished lecithotrophic larvae of *O. glabrum* would be capable of dispersing even under food-impoverished conditions. The current study examines ophiuroid reproductive biology over multiple localities in the CCZ concurrently for the first time, at sites characterised by differing productivity regimes. The reproductive biology of each species is thus discussed with reference to past evolutionary (habitat stability), contemporary (food supply) and future environmental drivers (potential impacts of deep-sea mining).

## 1 Introduction

The challenges of exploring remote deep-sea abyssal environments have, thus far, insulated the deep-sea benthos from the impacts of mineral resource extraction. However, the advent of the technological means to access and exploit the considerable mineral resources found in these environments (Ghosh and Mukhopadhyay, 2000), heralds imminent, unprecedented levels of disturbance in the deep-sea (Weaver et al., 2018). Fauna that have evolved under highly stable, food-limited environmental regimes are likely to be poorly adapted to large-scale disturbances that rapidly and/or irreversibly alter their environment (Stearns, 2000). The Clarion-Clipperton Fracture Zone (CCZ) in the tropical NE Pacific exemplifies this scenario, where the largest known global reserve of polymetallic nodules is located, formed over geological timescales in abyssal soft sediments characterized by very low sedimentation rates (Hein et al., 2013). Recent studies that have sought to describe the benthic fauna that typify the CCZ have also revealed extraordinarily high taxonomic diversity across representatives from all Domains of Life (e.g. Amon et al., 2016; De Smet et al., 2017; Shulse et al., 2017; Wilson, 2017; Hauquier et al., 2018; Goineau and Gooday, 2019; Brix et al., 2020; Christodoulou et al., 2020), making the CCZ of critical importance for biodiversity conservation. In many of these studies, data (e.g. species-abundance curves) strongly suggest that many – arguably most – species remain unaccounted for, with high species turnover over relatively short spatial scales even in groups that are brooders like Isopoda (e.g. Wilson, 2017, Brix et al., 2020). Remoteness of habitat and spatial variability in species composition of this sort present challenges for performing robust ecological studies, as evidenced by the scarcity of ecologically meaningful data available for even the most conspicuous ‘common’ epifaunal species in the CCZ (Danovaro et al., 2017), including many echinoderm species (Amon et al., 2016).

Members of the phylum Echinodermata represent some of the most biomass-dominant taxa found in deep-sea abyssal plains (Gage and Tyler, 1991). Echinoderms play a significant role in global marine carbon budget (Lebrato et al., 2010) and are abundant in many soft- and hard-substrate habitats globally (Gage and Tyler, 1991). The class Ophiuroidea are known to be particularly prevalent and diverse on bathyal slopes (e.g. O’Hara et al., 2008) but data concerning species diversity and abundances at abyssal depths are scarce by comparison (Stöhr et al., 2012). In the equatorial NE Pacific, such data is limited to select historic (e.g. HMS Challenger and Albatross expeditions, Lyman, 1878, 1879, 1882; Clark, 1911, 1949) and contemporary surveys (Amon et al., 2016; Vanreusel et al., 2016, Christodoulou et al., 2020). Drivers for large-scale regional variability in abyssal ophiuroid population densities are still poorly understood; however, in the CCZ it seems that local-scale patchiness in nodule substrate availability plays a key role in determining the distribution of many species. Under certain habitat conditions, ophiuroids and the echinoderms more generally, comprise one of the largest components of mobile epifauna at nodule-rich sites (Amon et al., 2016; Vanreusel et al., 2016: up to 15 individuals per 100 m^2^, with major contributions from ophiuroids). The presence of nodules appears particularly relevant to certain ophiuroid and echinoid species; when nodules are absent, mobile epifaunal densities fall sharply to one or two encounters in an equivalent 100 m^2^ area, largely due to much-reduced encounters with ophiuroid species (Vanreusel et al., 2016). Two recent studies published in 2020, one based on video-transect surveys and the other on specimens collected by remotely operated vehicle (ROV) and epibenthic sledge (EBS), have also identified ophiuroids in abundance and notably, at unexpectedly high levels of diversity in the easternmost regions of CCZ (Christodoulou et al., 2020; Simon-Lledó et al., 2020) with the discovery of previously unknown ancient lineages (Christodoulou et al., 2019, 2020).

Current knowledge of reproductive biology in abyssal ophiuroid species is limited to a few papers examining gametogenic and/or size-class patterns (e.g. *Ophiomusa lymani* in N Atlantic & NE Pacific, Gage, 1982; Gage and Tyler, 1982; *Ophiocten hastatum* in NE Atlantic, Gage et al., 2004; *Ophiura bathybia, Amphilepis patens, Amphiura carchara* and *Ophiacantha cosmica* in NE Pacific, Booth et al., 2008; *Ophiura irrorata loveni, Ophiura lienosa, Amphioplus daleus, Ophiacantha cosmica, Ophiernus quadrispinus* and *Ophioplexa condita* in S Indian Ocean, Billett et al., 2013), with no data available for specimens collected from polymetallic-nodule habitats. Two of the more frequently encountered ophiuroids within the eastern CCZ (Christodoulou et al., 2020) are the brittle stars *Ophiosphalma glabrum* Lütken & Mortensen, 1899 (Ophiosphalmidae) and *Ophiacantha cosmica* Lyman, 1878 (Ophiacanthidae). These two species make for an interesting comparative reproductive study due to their contrasting biology. *Ophiosphalma glabrum* is relatively large (35 – 40 mm maximum disc diameters, Clark, 1911, 1913), epifaunal on soft sediments (0 – 2 mm burial, Amon et al., 2016; Glover et al., 2016; Christodoulou et al., 2019) and likely a generalist deposit feeder, as is the case in the closely related genus *Ophiomusium* (Pearson and Gage, 1984). *Ophiacantha cosmica*, by contrast, is considerably smaller (11 – 12 mm maximum disc diameters, Booth et al., 2008; Billett et al., 2013) and epizoic or epifaunal on hard substrata (Billett et al., 2013), where it filters feeds with its spinous arms (Pearson and Gage, 1984). Although reproductive data are currently not available for *Ophiosphalma glabrum*, a few data already exist for *Ophiacantha cosmica* in nodule-free habitats. These indicate that *O. cosmica* is probably both gonochoric and lecithotrophic (oocyte 30 – 560 µm in diameter in specimens from the Southern Indian Ocean, Billet et al., 2013), with seasonal fluctuations in body-size structure suggesting that specimens spawn in response to peaks in particulate organic-carbon (POC) flux (Booth et al., 2008).

As part of a wider concerted effort to address significant knowledge gaps in our ecological understanding of nodule-rich seabeds while this habitat remains relatively pristine, the current study describes the reproductive biology of the brittle stars *Ophiosphalma glabrum* Lütken & Mortensen, 1899 (Ophiosphalmidae) and *Ophiacantha cosmica* Lyman, 1878 (Ophiacanthidae) in a nodule-rich environment, based on histological analyses of specimens collected from several mining-contract areas within the eastern CCZ.

## 2 Methodology

### 2.1 Sample collection, fixation, preservation and species identification

Specimens were collected during remotely operated vehicle transects (ROV Kiel 6000, GEOMAR, manipulator arm) undertaken over the course of two cruises in Spring 2015 and 2019 on the R/V Sonne in the eastern CCZ. Cruise SO239 (11 March – 30 April 2015) visited four contract areas and one Area of Particular Environmental Interest (APEI, 400 x 400 square-km protected areas assigned to each of nine presumptive ecological subregions of the CCZ). The 2015 samples in the current study originate from three of these contract areas where one or both of the species in the current study were found: BGR area (Federal Institute for Geosciences and Natural Resources, Germany); the southern IOM area (Interoceanmetal Joint Organization); and the easternmost GSR area (G-TEC Sea Mineral Resources NV, Belgium), where each was located along a decreasing SE-to-NW POC gradient, concomitant with a decrease in surface primary productivity driven by basin-scale thermocline shoaling towards the N. Pacific subtropical gyre (Pennington et al., 2006). The 2019 samples from the follow-up cruise SO268 (18 Feb – 21 May), originated from BGR (again) and the central GSR contract area (2015-2019 site separation: BGR ∼ 50 km; GSR ∼ 350 km). A site map is provided in Figure 1.

**Figure 1.**
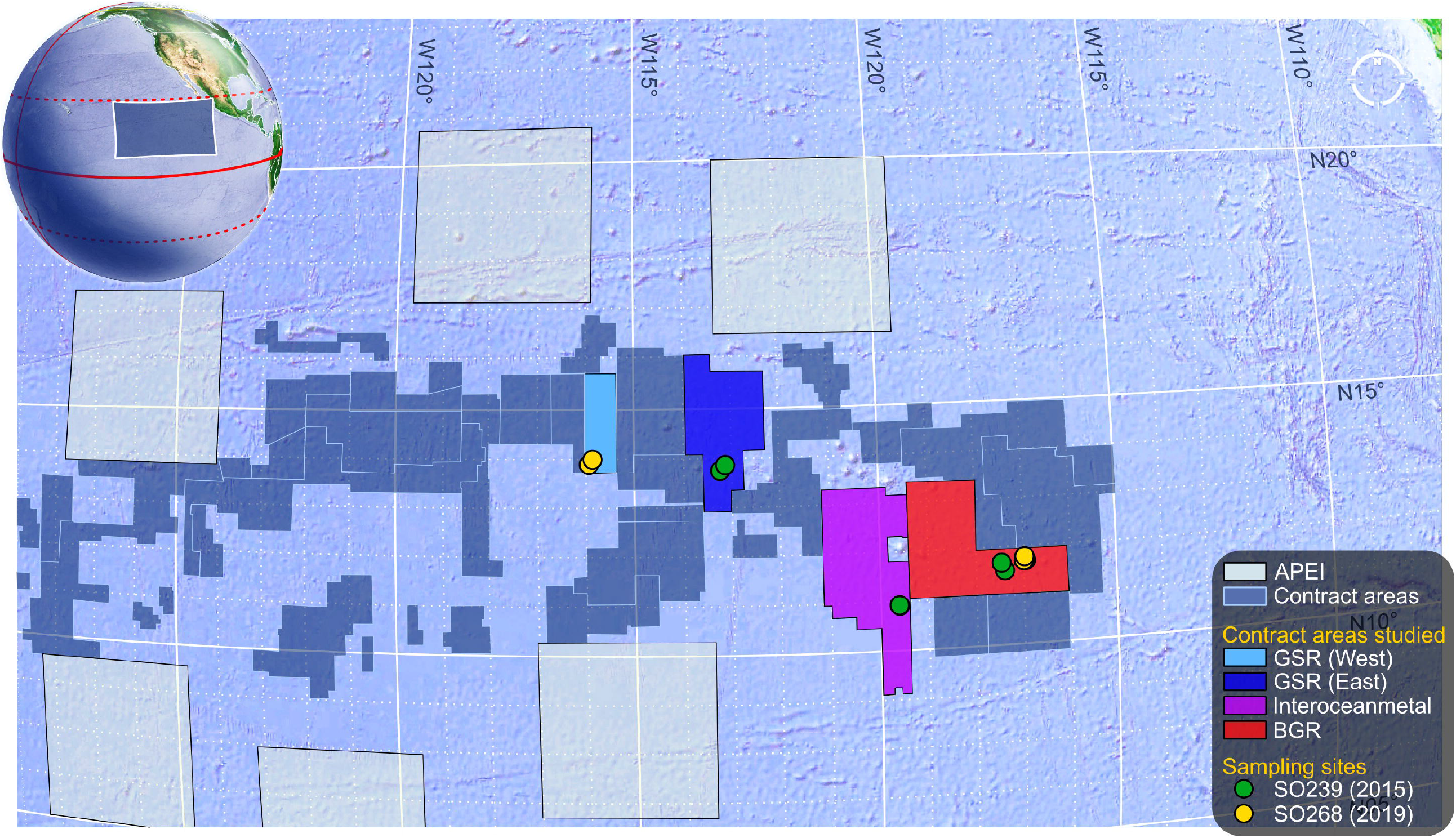
Map of eastern Clarion-Clipperton Fracture Zone (CCZ) with sampling details. Map depicts the eastern half of the Area within the CCZ. Large 400 x 400 km square regions are Areas of Particular Environmental Interest (APEI) that border numerous exploration contract areas in the centre (N.B. so-called “reserved” areas are not shown). The areas in which specimens were collected are identified along with site locations for each cruise. These were from the BGR area (Federal Institute for Geosciences and Natural Resources, Germany); the southern IOM area (Interoceanmetal Joint Organization); and both the central and the easternmost GSR areas (G-TEC Sea Mineral Resources NV, Belgium).

Following collection, the arms of each specimen were removed for molecular species identification, while the central disc was fixed for reproductive histology. Arm fixation was in pre-cooled 96% EtOH, with replacement after 24 h (stored at -20 °C thereafter); disc fixation was in 4% phosphate-buffered formaldehyde (48 h) following by serial transfer to 70% ethanol, stored at room temperature. Specimens were later identified to species level based on morphological characteristics (*Ophiosphalma glabrum*: Lütken and Mortensen, 1899; Baker, 2016; *Ophiacantha cosmica*: Lyman, 1878; Paterson, 1985), supported by mtDNA COI analyses of genomic DNA extracted from arm tissues, described in detail in Christodoulou et al. (2020).

### 2.2 Size measurements, dissection and tissue preparation for microscopy

Micrographs of the oral and aboral faces of central discs were created from focus-stacks taken under a stereomicroscope (Leica M125 and DMC5400 camera, LAS-Leica Application Suite 3.7; focus-stacking in Combine ZP 1.0). Disc diameter – the length from the radial shield’s distal edge to the opposing interradial margin (Gage and Tyler, 1982) – was measured in LAS, to estimate size at first maturity and assess size-class frequencies as a function of sex. The aboral face, stomach and bursa lining were then removed by dissection to reveal the bursal slits that border each arm base, along which the sac-like gonads are located, arranged in rows on each side. Intact gonad rows were dissected and then embedded either in paraffin wax or plastic resin, depending on tissue size. An additional decalcification step in 3% nitric acid in 70% ethanol (Wilkie, 2016) followed by a 70% ethanol rinse was required for smaller gonads, which were retained on a fragment of calcified tissue to facilitate tissue processing.

Larger gonad samples (all *Ophiosphalma glabrum* specimens and one *Ophiacantha cosmica* specimen with larger gonads) were infiltrated in molten paraffin at 60 °C, following 6-step serial transfer to 100% ethanol, 4-step transfer to 100% MileGREEN™ (an iso-paraffin based solvent, Milestone) and final 2-step transfer to paraffin in a mould to set. Blocks were trimmed and 5-µm sections cut, relaxed in a water bath, placed on Superfrost™+ slides and dry-fixed by incubation at 50 °C. Slides were stained in (Harris’) Haematoxylin & Eosin(-Y) following standard protocols with dehydration by replacement using ethanol. Note that the *O. cosmica* specimen processed using paraffin wax was also processed with LR white resin (protocol below) using neighbouring gonads, to identify any procedural biases in oocyte feret diameter.

Smaller tissue samples (all *O. cosmica* specimens) were blotted dry and then infiltrated directly in a hydrophilic methacrylate resin (LR white, London Resin Co.) at room temperature (3 resin replacements by micropipette, 30-mins. each, 4th overnight) then transferred to a fresh resin-filled gelatine capsule (size 00, Electron Microscopy Sciences, UK), which was capped and polymerised at 50 °C (20+ hrs). Having removed gelatine, LR-white resin pellets were trimmed and wet-sectioned on a Leica Ultracut UC6 Ultramicrotome (Germany), using a 45° glass knife. Sections 1-µm thick were transferred to individual aliquots arranged on Superfrost™+ slides. Periodic toluidine staining tracked progress through tissue. Standard H & E staining was modified by extending staining times and excluding ethanol, which distorts LR white. Slides were dried at 50 °C with a desiccant (see Laming et al., 2020 for details). Regardless of protocol, sections were then cleared in MileGREEN™ and mounted with a cover slip (Omnimount™, Electron Microscopy Sciences).

### 2.3 Gametogenic analyses

Stained serial sections were photographed under a compound microscope (Leica DM2500 with camera module ICC50W, LAS 3.7). Spermatogenesis was documented qualitatively only. Oogenesis was examined in more detail. Oocyte size frequencies were determined by image analysis of serial sections from ovaries, imported as image-stacks into Image-J (Schindelin et al., 2012) to track individual oocytes through the sample and prevent repeat counting. Up to 100 oocytes (with visible nuclei only) were counted per specimen, each measured along its longest axis (feret diameter) and classified into oogenic developmental stages, based on the presumption that germinal vesicle breakdown, GVBD, precedes oocyte release (Adams et al., 2019). Stages were: 1) *Previtellogenic* oocytes (Pv), smallest oocytes with no clear evidence of a germinal vesicle or vitellogenesis (i.e. eosinophilic granulated cytoplasm, rich in yolk proteins, has not yet developed); 2) *Vitellogenic I* (VI), “early” vitellogenic oocytes that are less than twice the diameter of previtellogenic oocytes, with the beginnings of an eosinophilic cytoplasm and a distinct germinal vesicle; 3) *Vitellogenic II* (VII), “mid-stage” vitellogenic oocytes that clearly possess an eosinophilic granulated cytoplasm, but whose diameter is less than double that of the germinal vesicle; 4) *Vitellogenic III* (VIII), “late” vitellogenic oocytes with large eosinophilic cytoplasmic reserves, whose diameter exceeds double that of the germinal vesicle; 5) *Asymmetric* (As), late vitellogenic oocytes with germinal vesicles located asymmetrically and in contact with the cell membrane, the precursor to germinal vesicle breakdown (GVBD) and onset of meiosis I; 6) *Polar bodies* (PB), oocytes that have undergone GVBD, with evidence of peripheral polar bodies, germinal vesicle lacking; and 7) *Mature* (M), fully mature oocytes, possessing only a small pronucleus (after Wessel et al., 2004).

Overall state of gonad maturity was assessed visually in terms of gonad aspect within the bursa and histological appearance. Gonad state was either defined as “immature” (no gonad), “no gametes” (gonad but no gametogenesis), “developing” (gametogenesis at all stages of development but with interstitial spaces remaining, equivalent to stages I and II in Tyler, 1977), “nearly ripe” (late-stage gametes dominate – spermatozoa in males or *VIII*-to-*Mature*, denoted here as “VIII_plus_” in females – but gonad is not yet replete, equivalent to stage III in Tyler, 1977), “ripe” (fully mature gametes and no interstitial space, equivalent to stage IV in Tyler, 1977) and “post-spawning / recovery” (some residual near-mature gametes remain alongside degraded material, epithelial and myoepithelial layers are thickened and nutritive phagocytes are typically present, equivalent to stages V in Tyler, 1977). For *Ophiosphalma glabrum*, the percentage area occupied by oocytes relative to gonad area and a maturity stage index (MSI) were also calculated, the latter using the 4^th^ formula proposed from Doyle, Hamel and Mercier (2012): *Oocyte density x Size of individual* ^*-1*^ *x Oocyte surface area x 0*.*01*, where oocyte density in the current study refers to oocytes mm^-2^ of all gonads in cross section, size of individual is disc diameter in mm (whose reciprocal value compensates for any size bias) and oocyte surface area is derived from the mean feret diameter (¼π*d*^2^).

### 2.4 Statistical analyses

Sex ratios for each species were assessed by χ^2^ test. The statistical significance of rank-based differences was tested by Kruskal-Wallis H test for: 1) disc diameter as a function of sex (including a juvenile category); 2) oocyte feret diameter as a function of year, and of disc diameter (for a given visual gonad state) and; 3) percentage gonad occupied by oocytes, MSI and mean oocyte feret diameter each as a function of visual gonad state. Post-hoc pairwise comparisons were by Dunn test. To assess whether a trade-off exists between oocyte diameter and oocyte density (and thus, fecundity), a Spearman rank-order correlation was used to assess the strength of any proportional relationship between oocyte density and mean oocyte feret diameter. Expressions of variation around average values are standard deviation (sd., for means) and median absolute deviation (mad., for medians). All statistical and graphical analyses were performed in R 4.0.3 (R Core Team, 2020)^1^, using packages *svglite, tidyverse, cowplot, ggExtra* and *ggpubr* (Attali and Baker, 2019; Wickham et al., 2019, 2020; Kassambara and Kassambara, 2020; Wilke, 2020), with figures prepared in Inkscape 1.0.2 (Inkscape Project)^2^.

## 3 Results

### 3.1 Sex ratios, size-class frequencies and size at first maturation

A total of seventy-six *Ophiosphalma glabrum* individuals (22 in 2015, 54 in 2019) and twenty-three *Ophiacantha cosmica* individuals (8 in 2015, 15 in 2019) were processed. Molecular analyses of the arms of specimens in the current study used for species’ confirmation in a subset of individuals have been compiled into dedicated DOI-indexed dataset containing accession codes (Genbank), BOLD IDs and photos, trace files and collection data and specimen metadata <u>(ht</u>t<u>p://dx.doi.org/10.5883/DS-</u> CCZ4). Of the two species, *Ophiosphalma glabrum* was larger (across entire sampling area, mean disc diameter 16.0 ± sd. 4.02 *vs. Ophiacantha cosmica* mean disc diameter 4.9 ± sd. 2.03, example specimens in Figure 2). Diameters ranged from 5.7 – 25.2 mm in *O. glabrum* and 2.2 – 10.0 mm in *Ophiacantha cosmica*. Although male *Ophiosphalma glabrum* specimens were numerically dominant (39 *vs*. 25 individuals), overall sex ratios did not differ significantly from a 1:1 ratio (χ^2^_1_ = 3.06, p > 0.05). The three intermediate size classes for *O. glabrum* (together ranging from 12.5 – 20 mm) were characterised by equal numbers of males and females; however, smaller and larger mature *O. glabrum* individuals outside these size classes were exclusively male (Figure 3). Ranked-disc diameters did not significantly differ between sexes in *O. glabrum*, only between mature (male/female) and immature specimens, resulting in significant global differences across the three categories when immature specimens were included in analyses (Kruskal-Wallis χ^2^_2_ = 31.39, p < 0.01, Figure 3). Sample sizes were particularly small for specimens of *Ophiacantha cosmica*. Despite this, due to an over-whelming dominance of females (7:1) sex ratios were found to be significantly different (χ^2^_1_ = 4.50, p < 0.05). The only male *O. cosmica* collected was the largest individual of this species, making statistical comparisons of disc diameter between sexes impossible.

**Figure 2.**
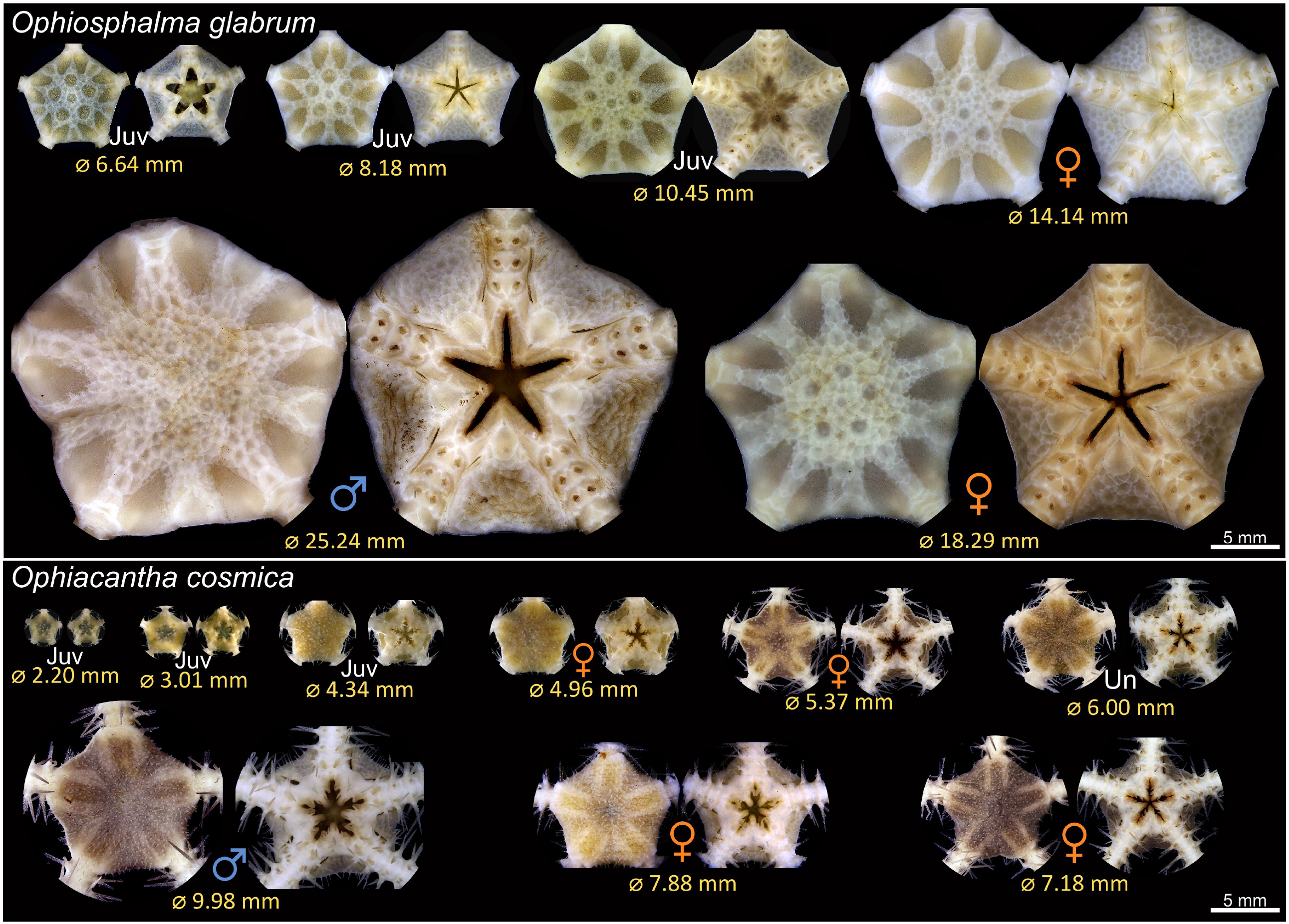
Example discs of *Ophiosphalma glabrum* and *Ophiacantha cosmica* from the current study. The aboral and oral faces of a subset of specimens encountered during the SO268 cruise in 2019 are pictured, representative of the size ranges encountered. Abbreviations: **Juv** *juvenile*; **Un** *undetermined sex* (see main text).

**Figure 3.**
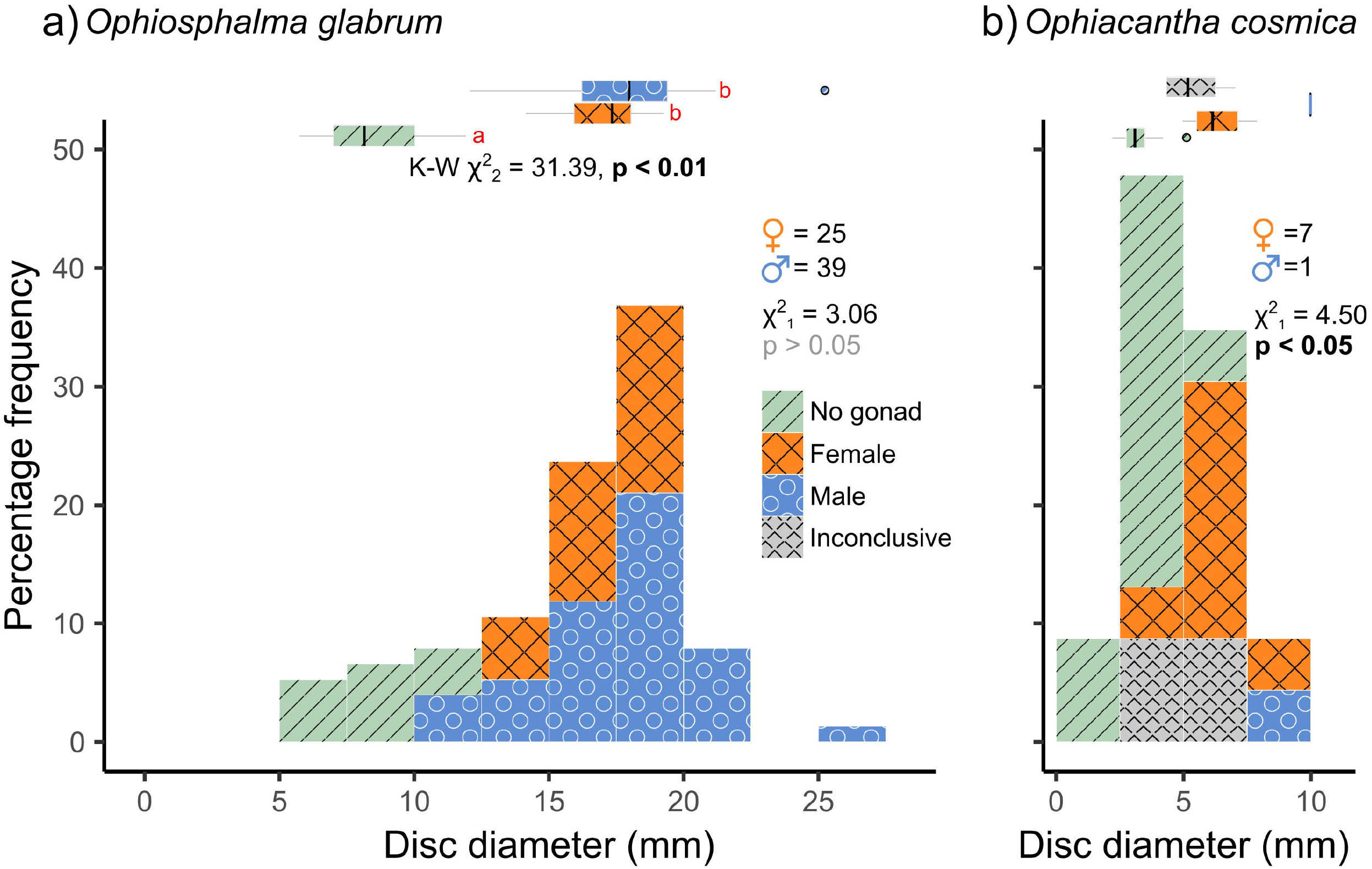
Size-class frequency distributions of *Ophiosphalma glabrum* and *Ophiacantha cosmica* disc diameters. Frequency histograms are colour-coded by sex (including a ‘No gonad’ category for immature specimens and an “Inconclusive” category for specimens with gonads devoid of gametes). Main χ^2^ statistics relate to test for deviation from a 1:1 sex ratio (M vs F only). Marginal box and whisker plots depict medians (vertical line), inter-quartile (box width) and 90% (whisker) ranges in disc diameter for each sex category. Proximate K-W χ^2^statistic and p-value in a) relate to Kruskal-Wallis test for ranked differences in disc diameter as a function of all sex categories in *Ophiosphalma glabrum*. Significant post-hoc pairwise comparisons (Dunn’s tests) are those with no letter annotations in common. Size-classes for each species are the same for comparative purposes.

For both species, several immature specimens were retrieved (Figures 2 and 3), allowing rough estimates of size at first maturity to be made. In *Ophiosphalma glabrum*, the largest immature specimen (lacking discernible gonads) had an 11.93-mm wide disc; a further 11 smaller specimens also lacked gonads. The smallest *O. glabrum* individual that possessed a gonad was a male at 12.07 mm disc diameter, while the smallest female was 14.14 mm (gametogenesis observed in both), placing approximate sizes at first maturity in males at < 12 mm disc diameter and at < 14.2 mm disc diameter in females. In *Ophiacantha cosmica*, size at first maturity is less concrete. The largest specimen with no discernible gonad had a disc diameter of 5.12 mm, with two smaller (4.28, 4.34 mm) and two larger (6.00, 7.04 mm) specimens each in possession of 1 – 2 very small gonad buds (ca. 100µm diameter) not yet furnished with gametes. The smallest individual in which gametogenesis was identified was a female of 4.96 mm with six larger females with disc diameters of 5.37 – 7.88 mm (Figure 3, select specimens in Figure 2). The largest individual – the only *O. cosmica* male – had a 9.98-mm wide disc.

### 3.2 Gonad state and gametogenesis

Gonad state and gametogenesis were assessed qualitatively in males (Figure 4 and 5) and both qualitatively and semi-quantitatively in females (Figures 4, 5 and 6). No evidence of simultaneous or sequential hermaphroditism (i.e., ovotestes / sex-specific size stratification) were identified in either species. Most *Ophiosphalma glabrum* males’ testes were *developing* (49 %) with easily identifiable spermatogenic columns, composed of progressively more-advanced stages of spermatogenesis, extending from the inner-sac membrane into the central lumen (Figure 4a). Testes were somewhat angular (Figure 4g), lacking the creamy engorged appearance typical of ripe individuals (example in Figure 4h). A further 35 % were considered *nearly ripe*, where testes aspect appears unchanged but spermatogenic columns are no longer clearly visible, with roughly equal proportions of central mature spermatozoa, and surrounding tightly packed cells at earlier stages of spermatogenesis. Three specimens were *ripe* (8 %), where gonads appear engorged, extensively spread throughout bursae and replete with spermatozoa (Figure 4b, h). Putative evidence of nutritive phagocytes was used as the basis for discriminating a minority of male specimens as being in a *post-spawn / recovery* state (8 %, Figure 4 c, with gonad aspect in i). The single *Ophiacantha cosmica* male collected was at the *developing* stage, with evident spermatogenic columns (Figure 5a).

**Figure 4.**
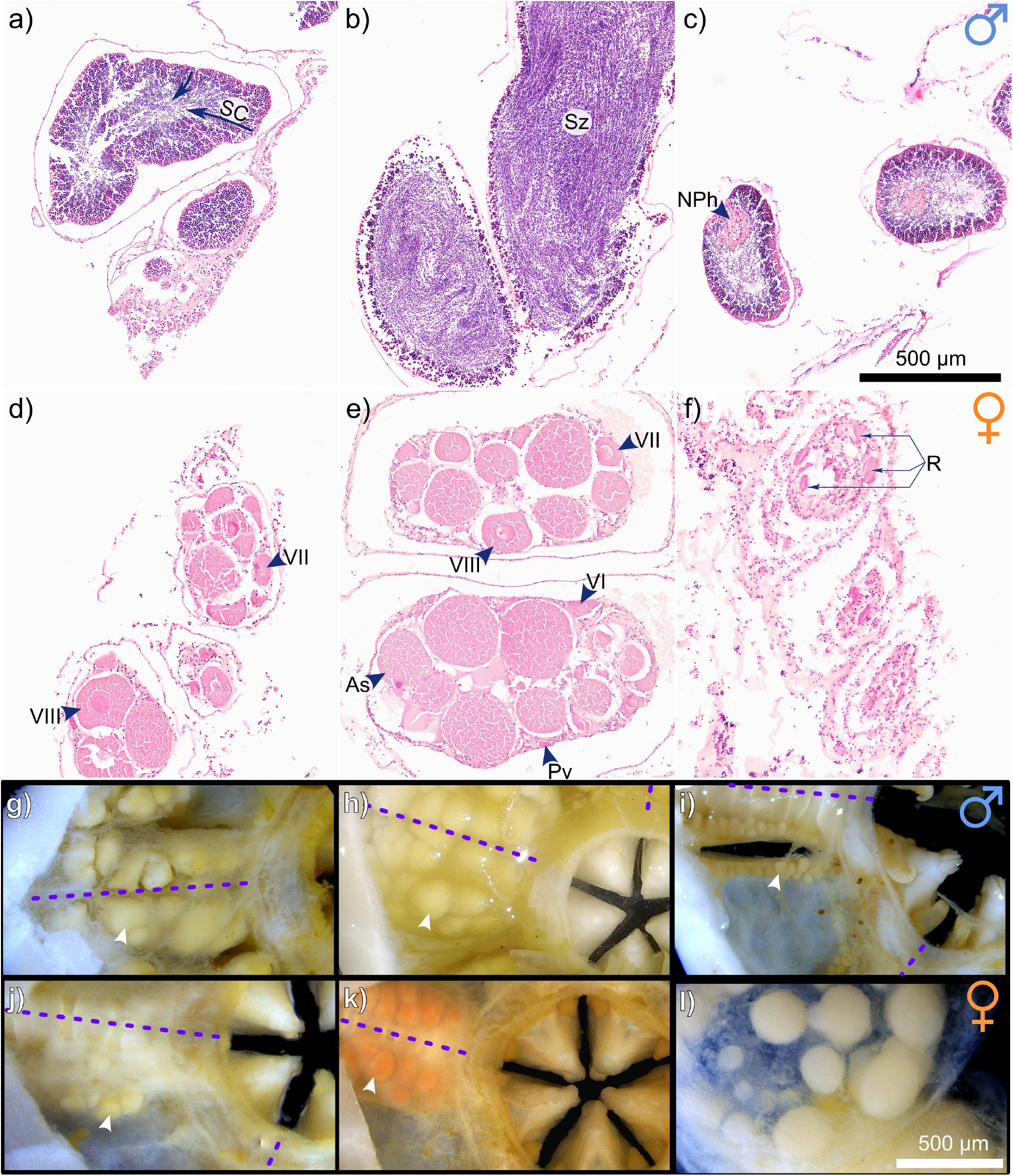
Gametogenesis and gonad development in *Ophiosphalma glabrum*. Gonads at various stages of development in *O. glabrum*, as follows: a) *developing* testes in which spermatogenic columns (**SC**) are evident (indicated with arrows), b) *ripe* testes and c) *post-spawn* / *recovery* testes; d) *developing* ovaries, e) *nearly ripe* ovaries and f) *post-spawn* / *recovery* ovaries; visual aspect of g) *developing* testes [corresponding to a], h) *ripe* testes [corresponding to b] and i) *post-spawn* / *recovery* testes [corresponding to c] and finally, visual aspect of j) *developing* ovaries [corresponding to d], k) *nearly ripe* ovaries [corresponding to e] and l) a single ovary viewed down the distal-proximal axis, where late-stage oocytes are located distally (foreground), having previously developed from the basal region where early-stage oocytes are visible beneath. White arrowheads in g-l indicate single gonads, purple dotted lines are midlines of arm bases. Other abbreviations: **As** *Asymmetric oocytes*, **NPh** *Nutritive phagocytes*, **Pv** *Previtellogenic oocytes*, **R** *Residual oocytes*, **VI** *Vitellogenic I oocytes*, **VII** *Vitellogenic II oocytes*, **Sz** *Spermatazoa*, **VIII** *Vitellogenic III oocytes*.

**Figure 5.**
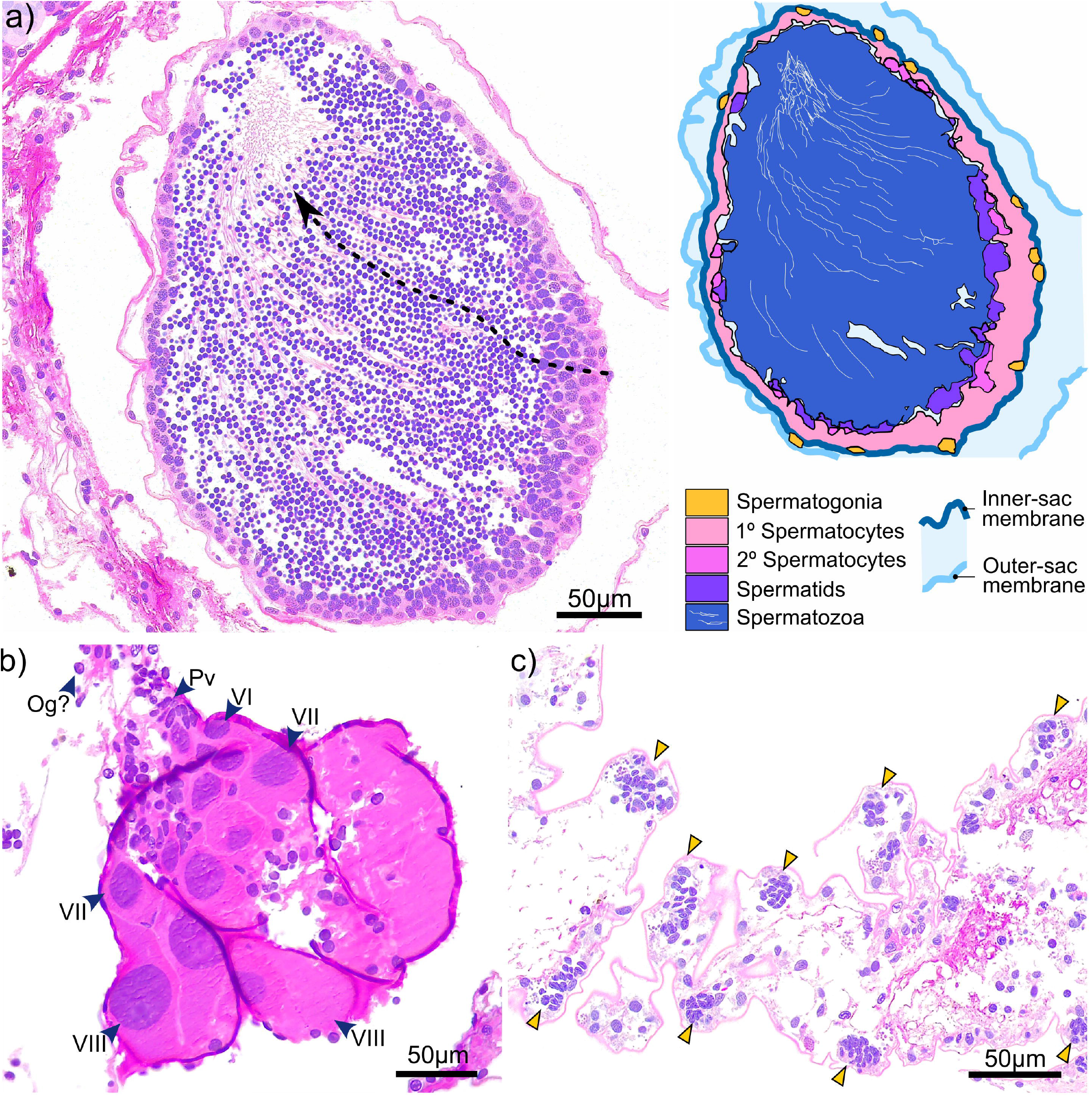
Gametogenesis and gonad development in *Ophiacantha cosmica*. Gonads at different stages of development in *O. cosmica*: a) *developing* testes, in which spermatogenic columns are evident (dotted arrow), where cells at progressive developmental stages form chains that extend from the inner-sac membrane into the central lumen; b) *developing* ovaries, in which various oogenic stages co-occur (largest oocytes are distal-most to region of gonad attachment); and c) pre-gametogenic gonadal tissue, with densely arranged pockets of putative primordial germ cells with basophilic nuclei that appear to form node-like gonadal precursors. Abbreviations: **Og** *Oogonia (putative)*, **Pv** *Previtellogenic oocytes*, **VI** *Vitellogenic I oocytes*, **VII** *Vitellogenic II oocytes*, **VIII** *Vitellogenic III oocytes*.

**Figure 6.**
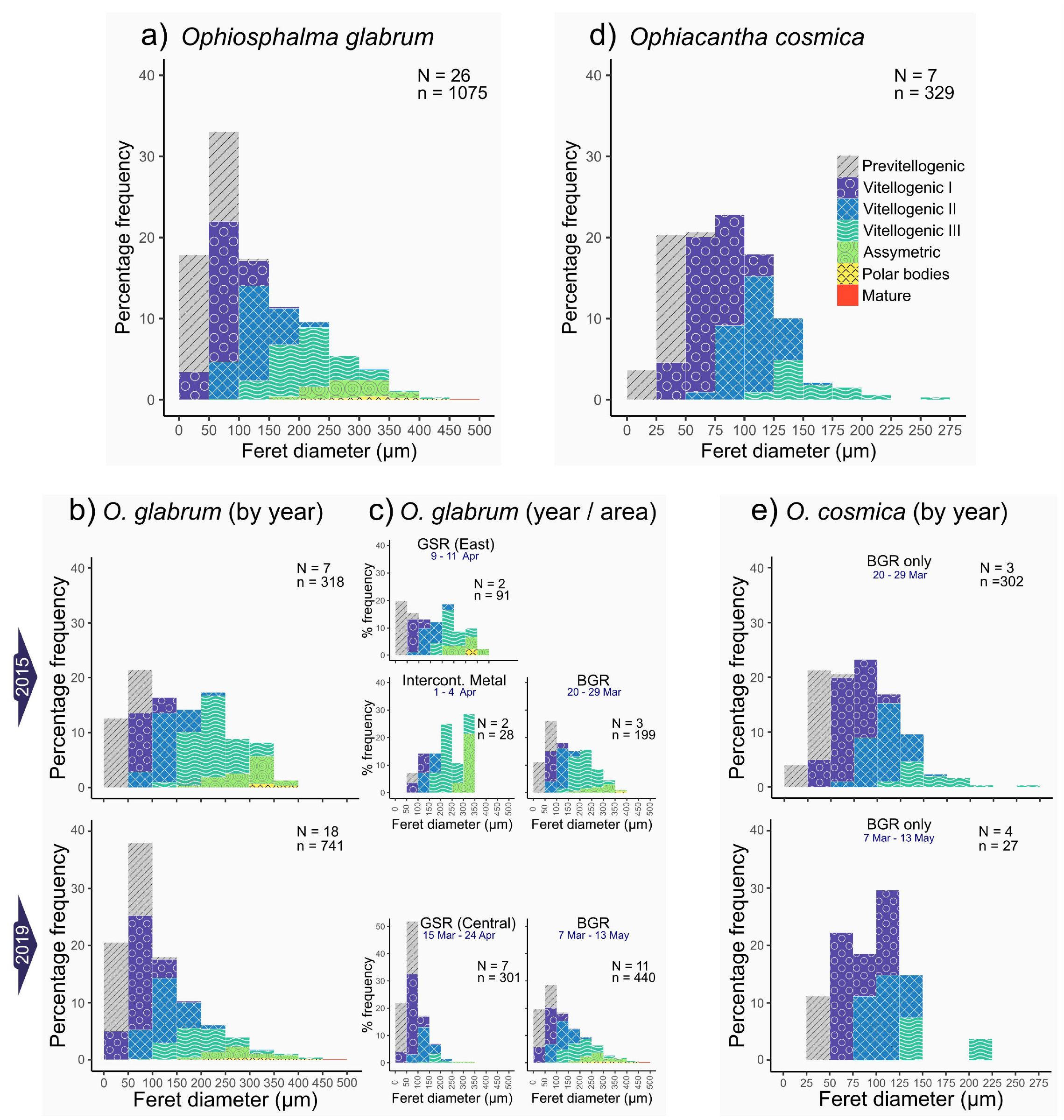
Oocyte size-class frequency distributions. Frequency histograms display the relative proportions of oocytes in each size-class for feret diameters measured in females of each species. Only oocytes with nuclei were considered. Colour coding relates to associated oogenic developmental stages, assessed on a case-by-case basis for each oocyte counted and measured. Data for *Ophiosphalma glabrum* are on the left (a – c) displaying; a) whole *O. glabrum* dataset; b) data by sampling year and; c) data by site for each year. Data for *Ophiacantha cosmica* are on the right (d – e), displaying; d) whole *O. cosmica* dataset and e) data by sampling year (this species was only recovered from the BGR contract area). N = total number of females, n= total number of oocytes pooled from N females.

No *ripe* females were identified in either species. However, most *Ophiosphalma glabrum* females had ovaries in a *nearly ripe* state (56 % overall, 100 % in 2015, 39 % in 2019), where VIII_plus_ oocytes dominate but the ovary is not replete and has not yet spread far into the bursa (e.g. Figure 4e, k). A further 33 % of individuals in *post-spawn / recovery* and 27 % with *developing* ovaries (see Figure 4d and f) were also identified, all in 2019. All-but-one of the *Ophiacantha cosmica* females were *developing* (e.g., Figure 5b). The smallest female was in a *post-spawn / recovery* state (or possibly *developing* for the first time).

Oocytes at every stage of oogenesis were identified in *Ophiosphalma glabrum* (Figure 6a, b and c). Maximum oocyte feret diameters for *Ophiosphalma glabrum* and *Ophiacantha cosmica* were a mature oocyte of 453 µm and VIII oocyte of 273 µm, respectively. However, unlike *Ophiosphalma glabrum*, oocytes at stages more advanced than VIII were not observed in *Ophiacantha cosmica* generally (Figure 6d and e), so it is likely that maximum oocyte diameters in this species are underestimated. The higher proportion of *nearly ripe Ophiosphalma glabrum* females in 2015 versus 2019, is reflected in significant differences in ranked oocyte feret diameters between 2015 and 2019 (Kruskal-Wallis χ^2^_1_ = 63.5, p < 0.0001). These differences, echoed by mean and median oocyte ferret diameters for each year (2015-mean 159.1 ± sd. 91.7, median 149.5 ± mad. 109.0; and 2019-mean 112.0 ± sd. 78.3, median 88.0 ± mad. 57.8), relate to higher frequencies of VIII_plus_ oocytes in the ovaries of specimens from 2015, clearly visible in the size-frequency data for both years (Figure 6b). Unfortunately, low sample sizes prohibit the quantitative comparison of oocyte frequency distributions as a function of contract area. That said, the proportion of large, late-stage oocytes found in specimens at the eastern-most sites appears higher (particularly for 2019, Figure 5c). No significant differences in ranked oocyte feret diameters between the two sampling years was found for *Ophiacantha cosmica* (Kruskal-Wallis χ^2^_1_ = 2.34, p > 0.05), with similar oocyte size-class means, medians and distribution profiles identified for this species in both years (2015-mean 82.3 ± sd. 40.7, median 79.1 ± mad. 43.7; and 2019-mean 94.4 ± sd. 40.0, median 90.8 ± mad. 39.3; profiles in Figure 6e), though very low sampling sizes are likely undermining the statistical power of the test.

In addition to classifying oogenic stage and measuring oocyte feret diameters, the cross-sectional area occupied by oocytes relative to gonad area (inner-sac membrane) was calculated for *Ophiosphalma glabrum* – where tissue section quality allowed – as a proxy for the degree to which female gonads were full. A maturity stage index (MSI) previously demonstrated to be more sensitive to subtle changes in gonad maturation (Doyle et al., 2012), was also calculated for *O. glabrum*. Percentage of gonad occupied by oocytes (Kruskal-Wallis χ^2^_2_ = 11.37, p < 0.01), mean oocyte feret diameter (Kruskal-Wallis χ^2^_2_ = 10.55, p < 0.01) and MSI (Kruskal-Wallis χ^2^_2_ = 9.13, p < 0.05) all differed significantly as a function of *O. glabrum* gonad state (Figure 7). Evident differences existed in MSI and in the relative gonad area occupied by oocytes in *nearly ripe* females; however, these metrics both failed to resolve differences between *developing* and *post-spawn* females, with post-hoc pairwise comparisons revealing significant group differences only between *nearly ripe* females and the two other gonad states (Figure 7). Group differences in mean oocyte feret diameter were restricted to *nearly ripe* females and those classified as *post-spawn / recovery* (Figure 6), reflecting similar (non-significant) mean feret diameters between *developing* and *nearly ripe O. glabrum* females. Spearman-rank correlative analysis revealed oocyte density mm^-2^ of gonad cross-sectional area to be significantly, inversely related to mean oocyte feret diameter (coefficient *R* = -0.74, p < 0.0001, Figure 7) indicating a trade-off between investment per oocyte (i.e., size of yolk reserves) and the space available to house oocytes in each gonad.

**Figure 7.**
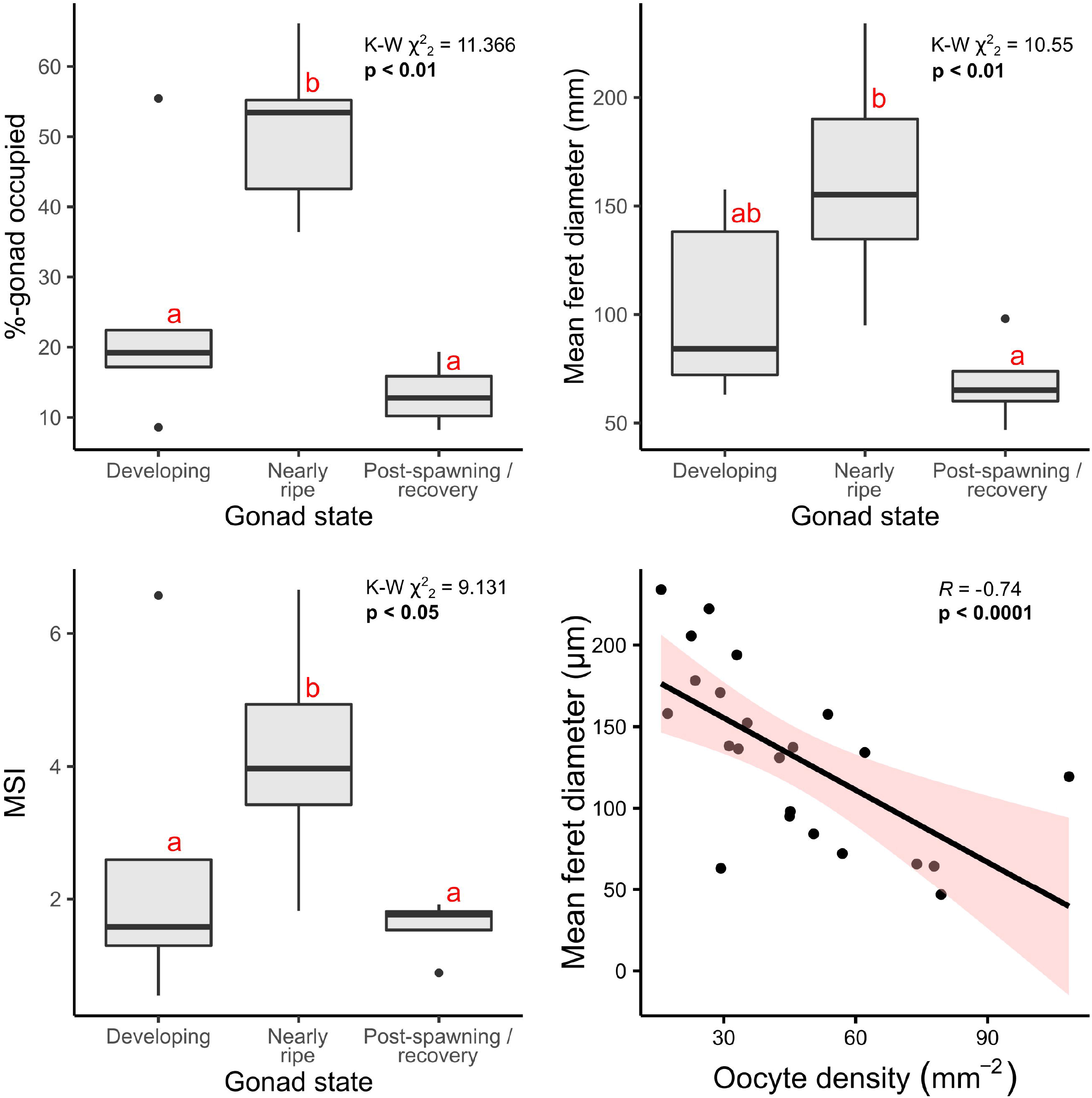
Reproductive metrics plotted as a function of gonad state and oocyte density. Analyses are based on 23 female specimens, of which 14 were *nearly ripe*, 5 were *developing* and 4 were in *post-spawning / recovery*. a) Percentage-gonad occupied by oocytes, b) overall mean ferret diameter per individual and c) a maturity stage index (MSI) developed by Doyle, Hamel, and Mercier (2012) were each calculated and evaluated as a function of gonad state, using Kruskall-Wallis Chi-squared tests on ranked data, the results of which are included in the top right corners for each. Significant post-hoc pairwise comparisons (Dunn’s tests) are those with no letter annotations in common. Depicted in d) is the Spearman rank-correlation analysis performed to identify any relationship between mean oocyte diameters and corresponding densities, with the resulting Rho statistic (R), p-value and 95-% confidence intervals displayed.

Finally, intraindividual variation in the number and size of ovaries and oocyte densities per ovary in *O. glabrum* was considerable. Several *developing* females possessed neighbouring gonads at different stages of maturity based on visual aspect (e.g. Figure 4j). The presence of oocytes at various stages of oogenesis resulted in large variability in oocyte diameters (Figure 8), with large variances around median values (Supplementary figure 1). Not unexpectedly – since one directly informed the other – the proportion of oocytes at each stage of oogenesis was relatively consistent in a given gonad state, irrespective of specimen size (Figure 8). However, ranked oocyte feret diameters overall were significantly higher in the largest *nearly ripe* disc-diameter size class, 17.5 – <20 mm, when compared with smaller size classes 12.5 – <15 mm and 15 – <17.5 mm (based on post-hoc comparisons, following a highly significant Kruskal-Wallis test for gonad state: Kruskal-Wallis χ^2^_2_ = 18.94, p < 0.0001, Supplementary figure 1). Significant post-hoc differences in ranked oocyte feret diameters were also identified across size classes in individuals in a state of *post-spawn / recovery* (Kruskal-Wallis χ^2^_2_ = 12.63, p < 0.01), due to the relative dominance of previtellogenic oocytes in one individual from the intermediate 15 – <17.5 mm size category for this gonad state (Figure 8). No significant size-related differences were identified in the oocyte feret diameters of *developing* individuals of either species.

**Figure 8.**
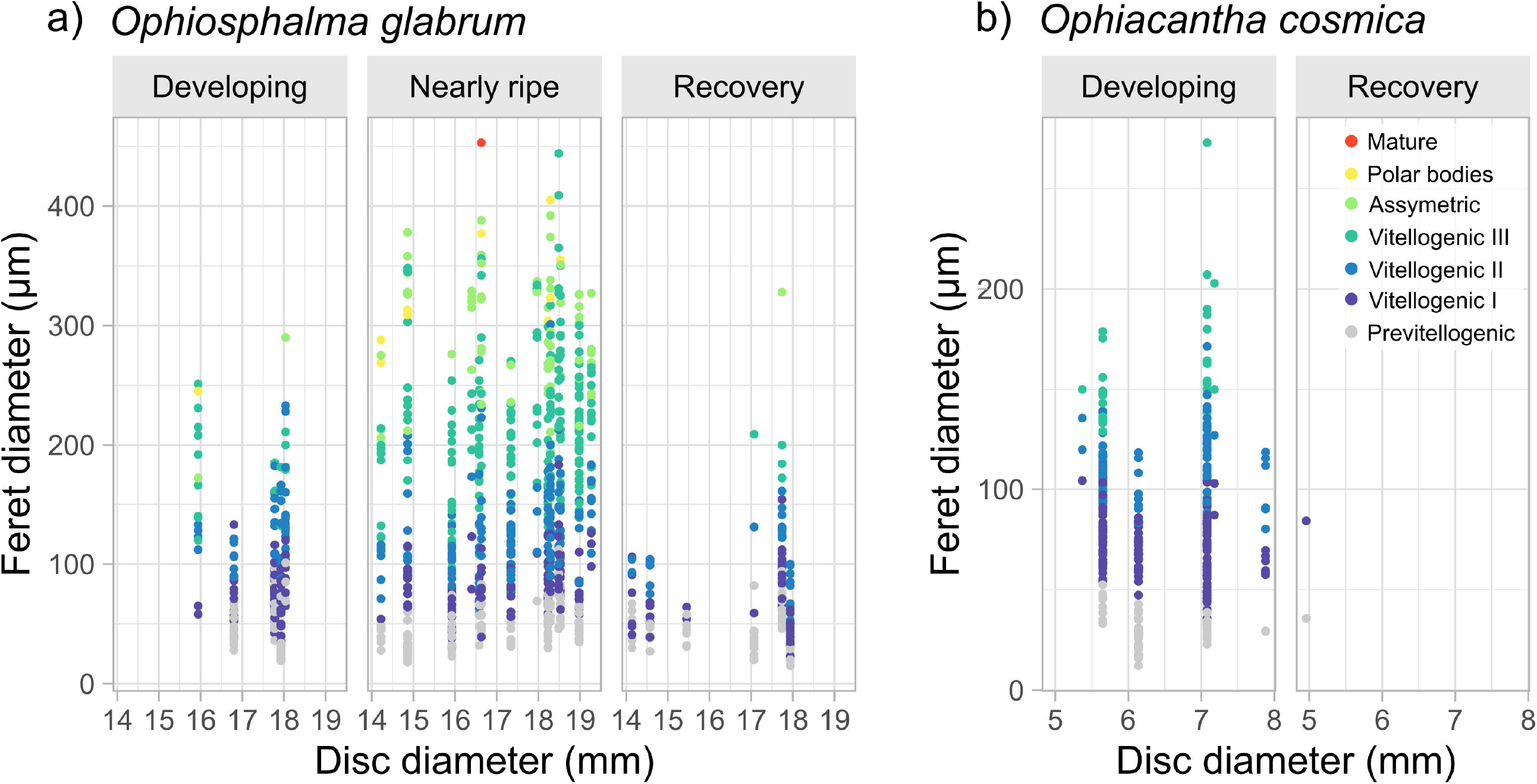
Variation in feret diameter in *O. glabrum* and *O. cosmica* with specimen size. Analyses are based on twenty-five *O. glabrum* specimens, of which 14 were *nearly ripe*, 5 were *developing* and 6 were in *post-spawning / recovery* and seven *O. cosmica* specimens, of which all but one was *developing*. Scatterplots display variation in oocyte feret diameters as a function disc diameter for female specimens in each gonad state identified, where colour-coding depicts oogenic developmental stage of each oocyte. Significant differences in oocyte feret diameter across specimen size classes were assessed for each gonad state (where possible), using Kruskall-Wallis Chi-squared tests on ranked data (see main text and Supplementary figure 1 for Box and whisker plots of binned disc-diameter data).

**Figure 9.**
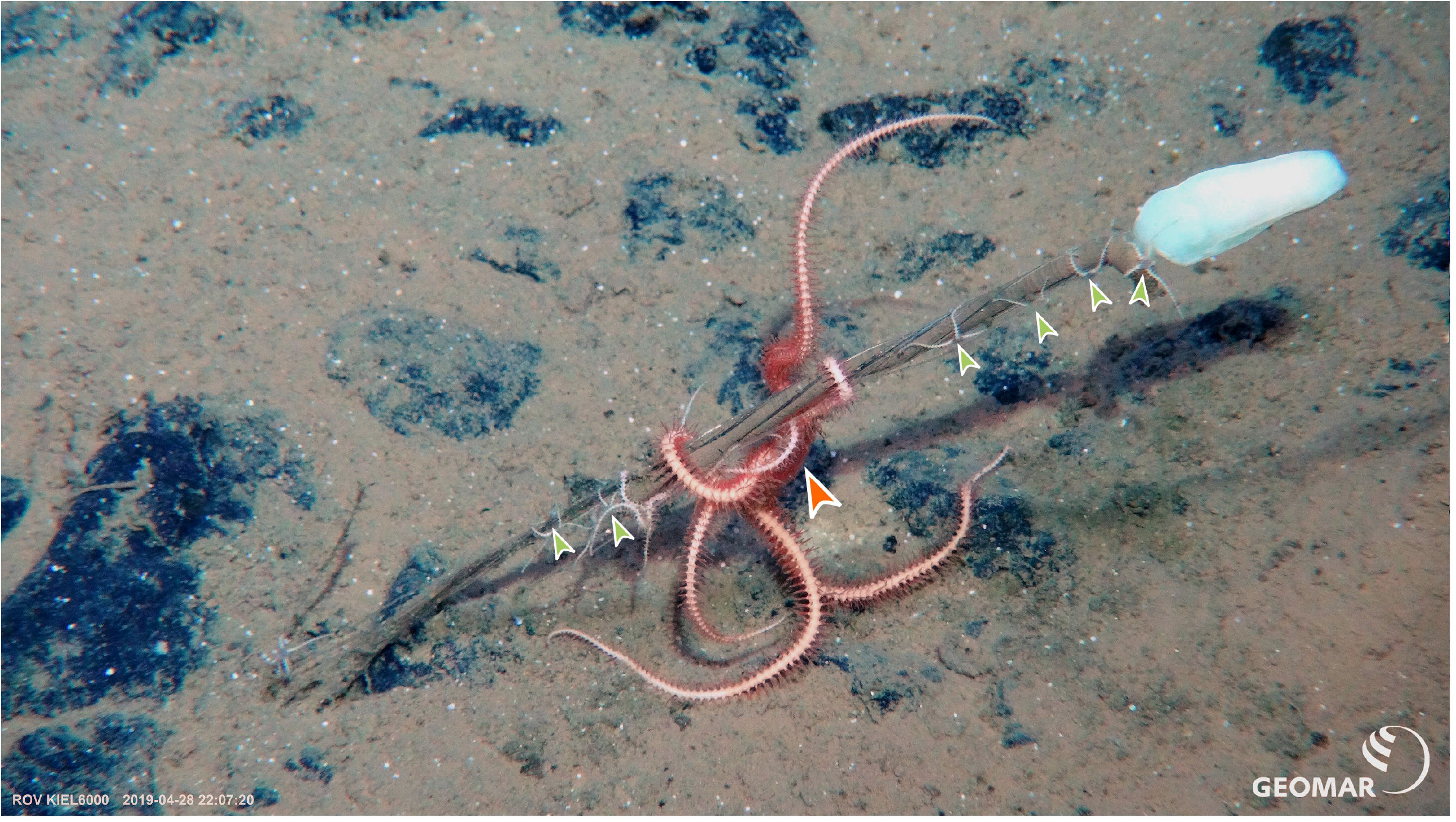
In-situ photograph of a group of *O. cosmica* on a stalked sponge. Pictured are two females (large orange arrowhead) and several juvenile specimens (small green arrowheads) of *O. cosmica* that were sampled in the BGR contract area in 2019 during the SO268 R/V Sonne cruise. NB. Dark rock-like deposits in sediment are polymetallic nodules. Photo credit: ROV Kiel 6000, GEOMAR

As a result of intra- and interindividual variability in gonad maturity overall, a reliable measure of fecundity proved impossible. However, extrapolating from 25µm of tissue sectioned per individual, late-stage (VIII_plus_) oocyte counts mm-sectioned^-1^ ovary ^-1^ ranged from 1.3 – 12.0 oocytes in *developing* females (n = 4), 5.0 – 30.0 oocytes mm-sectioned^-1^ ovary ^-1^ in *nearly ripe* females (n = 14) and 1.5 – 5.0 oocytes mm-sectioned^-1^ ovary ^-1^ in *post-spawn / recovery* females (n = 2).

## 4 Discussion

The current study provides new ecologically relevant reproductive data for two species of ophiuroid living in an oligotrophic, highly stable low-energy environment in the NE equatorial abyss which, due to the presence of polymetallic nodules, is under future threat from deep-sea mining activities (Weaver et al., 2018). The current study documents the reproductive biology of *Ophiosphalma glabrum* for the first time. Equal sex ratios for intermediate modal disc-diameter size classes and the absence of any simultaneous hermaphrodites both point to gonochorism in *O. glabrum*. Unfortunately, the manner (and possibly timing) of sampling may have created biases in the size classes of each sex. Large (and ripe) females (i.e., > 20-mm disc diameters) were conspicuously absent, suggesting either an unlikely upper limit on female disc diameter, or most probably, that large females were overlooked. This may simply be symptomatic of a patchy (or sex-specific) species distributions, where large communal echinoderm aggregations in the soft-sediment abyss only really occur during the episodic arrival of labile organic matter to the seafloor (Kuhnz et al., 2014; Smith et al., 2018) or during aggregative spawning events (Mercier and Hamel, 2009).

Oocyte feret diameters frequently exceeding 300 µm in the current study (max 453 µm in a mature oocyte) are indicative of a lecithotrophic (non-feeding) larval mode with substantial yolk-protein and lipid reserves (Sewell and Young, 1997; Young, 2003). Although this places an upper limit on transport time relative to planktotrophy, lecithotrophy releases larvae of nutritional constraints that likely exist in food-impoverished transport environments. In addition, if lecithotrophic dispersal occurs in deep waters where temperatures remain relatively low, this can act to extend transport times through metabolic suppression, with capacities for dispersal equivalent to planktotrophy (Mercier et al., 2013). Planktotrophic (feeding) larvae, by contrast, would be confined to water masses with higher detrital input (Young et al., 1997). Lecithotrophy necessitates greater energy investment per oocyte than in planktotrophy. This likely places constraints on fecundity in a food-limiting environment (Ramirez Llodra, 2002), as evidenced by the inverse relationship between oocyte densities and feret diameter in the current study. Ovaries in the current study, when *nearly ripe*, were classified as such based mainly on oogenic characteristics, since striking external indications of maturity were lacking in female specimens. Coloration was inconsistent and ovaries were not engorged, nor had they spread extensively into the bursae, as was witnessed in at least one ripe male. This indicates considerable scope remained for further reproductive investment in the females collected, under suitable environmental conditions. Without ripe females and intra- and interannual time-series sampling it is not possible to ascertain whether gametogenesis is seasonal (Tyler and Gage, 1980; Gage and Tyler, 1982) or aperiodic (‘continuous’), though the presence of all oogenic stages in most specimens suggests aperiodic or semi-continuous spawning behaviour (Brogger et al., 2013). Oocyte size-class data also appear to suggest that slight differences exist between 2015 and 2019, with a higher proportion of females approaching maturity in 2015. This could reflect the slightly different timing of cruises, as no *post-spawn recovery* females were found in 2015 or site-related differences for each year since site locations were not the same. Further data are necessary to confirm this.

In an oligotrophic environment lacking strong seasonal fluctuations in POC flux, apparent aperiodic (or ‘continuous’) gametogenesis may arise in species that spawn 1) periodically, but histological studies fail to identify their cyclical nature (Mercier et al., 2007; Mercier and Hamel, 2008) or 2) opportunistically, occurring in rapid response to increased food supply related to episodic massive changes in surface productivity and as such, remain undetected (Mercier and Hamel, 2009). Coupled with lecithotrophy, aperiodic opportunistic spawning would be resource-driven, allowing a species to respond to fluctuating food availability by investing in gametogenesis only when suitable conditions arise (e.g., Booth et al., 2008). Responsive reproductive modes of this sort appear to be a feature of bathyal and abyssal ecosystems in the tropical NE Pacific (Kuhnz et al., 2014; Smith et al., 2018) where the influences of episodic disturbance events are of greater influence than seasonal changes in surface productivity. However, additional sampling from highly oligotrophic NW regions of the CCZ (e.g. APEI 3, Vanreusel et al., 2016; Christodoulou et al., 2020) would be needed to establish whether lower POC flux translates into reduced energetic investment in reproduction.

Maximum sizes recorded for *O. glabrum* vary considerably in the literature but the largest attributed to this species is a 35 mm disc diameter (Clark, 1913), though there exists an *Ophiomusium multispinum* specimen – now synonymised with *O. glabrum –* with a 40-mm wide disc (Clark, 1911). In either case, the largest individual in the current study falls considerably short of these sizes, at 25.2 mm. Aside from some concerns around species assignation in historical samples and difficulties in collecting representative samples of species that occur at low densities in expansive and remote environments, these larger specimens also originate from shallower habitats (e.g. Cocos, Malpelo and Carnegie Ridge, Panama Basin, depths 800 – 3200 m, Clark, 1911, 1913), where greater food availability arriving from productive coastal systems (Pennington et al., 2006) could enable larger maximum sizes. Although smaller specimens were undoubtedly underrepresented, enough immature *O. glabrum* were collected to establish a rough size at first maturity of < 12 mm, around 30 - 35 % maximum size. Without knowledge of growth rates, the time taken to reach these sizes remains unknown; however, if measures of growth (e.g. Gage et al., 2004) or temporal size-class frequency analyses can be compiled in future studies, this value could be easily converted to an estimate of age at first maturity, a highly relevant conservation metric for response to disturbance.

Sample sizes for the smaller species *Ophiacantha cosmica* limit any detailed interpretation of their reproductive biology. However, some data on size-class distributions and oocyte feret diameters already exist for this species from nodule-free habitats at Station M in the NE Pacific and around Crozet Island in the Southern Indian Ocean (Booth et al., 2008; Billett et al., 2013). Although sex ratios were significantly different in the current study, little stock can be placed in this result as the total number of mature specimens numbered less than ten and maximum disc diameters for both sexes are markedly lower than in the literature. Evidence from other studies indicate that sex ratios follow a 1:1 ratio when sampled in larger numbers (Billett et al., 2013), with maximum disc diameters of 11 – 12 mm (Booth et al., 2008; Billett et al., 2013; M. Christodoulou, personal observations), although the disc diameter for this species’ holotype is unaccountably large, at 18 mm (Lyman, 1878). Maximum oocyte feret diameter in the current study (VIII oocyte at 273 µm) also appears to be underestimated, when compared with those of specimens from the Crozet Plateau (Billett et al., 2013), which reached maximum feret diameters exceeding 500 µm, indicating lecithotrophy (or direct development, though no evidence of brooding has ever been recorded, Billett et al., 2013; this study). This is to be expected as females in the current study were relatively small by comparison and only in *developing* states, with no oocytes in the final stages of meiosis (i.e., *asymmetric* or later). The only individual classified as *post-spawn* / *recovery* was probably

developing for the first time, particularly in light of its small disc diameter. Unlike the protandric hermaphrodite *Ophiacantha fraterna* (Tyler and Gage, 1982; species identification updated from *Ophiacantha bidentata* by Martynov and Litvinova, 2008), data from the current study – corroborated by that of Billett et al. (2013) – indicate that *O. cosmica* is gonochoric. However, the current study lacks sufficient data to confirm whether *O. cosmica* is also iteroparous, as is the case for *Ophiacantha fraterna* (Tyler and Gage, 1982). Size at first maturity for *O. cosmica* was assessed in the current study but remains approximate at < 4.96-mm disc diameter (in females), due to ambiguity around two relatively large specimens (6 – 7 µm disc diameters) that possessed tiny bud-like gonads in which no evidence for gametogenesis could be found. These gonads did possess nodes in which cell clumps with highly basophilic nuclei were located, which are believed to be gonadal precursors housing primordial germ cells (Figure 5c).

*Ophiacantha* spp. have been recorded from various hard substrata and vertically elevated fauna, such as deep-sea corals (Tyler and Gage, 1980) and tube worms (Lauerman et al., 1996; Billett et al., 2013). In the current study, this species is recorded on stalked-sponge species *Caulophacus* sp. and *Hyalonema* spp for the first time. Although alternative hard substrates are available in the CCZ (most obviously, polymetallic nodules), *O cosmica* specimens here were exclusively epizoic; this species was not encountered elsewhere. Stalked sponges are some of the most elevated of fauna on the seafloor in the CCZ; colonising them at positions above the benthic boundary layer would mitigate the influences of shear on current speeds, which likely aids in a suspension-feeding lifestyle (Booth et al., 2008). The specimens in the current study either occurred as single individuals or, in certain instances, as size-stratified groups (Supplementary figure 3), suggestive of gregarious behaviour, offspring retention or fissiparity (though pentameral *Ophiacantha* spp. are not known to be fissiparous, Lee et al., 2019). A more detailed molecular work, beyond the scope of this study, would help to reveal common or distinct genetic origins for the members of these groups.

In contrast, *Ophiosphalma glabrum* examined in the current study were epibenthic as found in previous studies both within the CCZ (Amon et al., 2016; Glover et al., 2016; Christodoulou et al., 2020) and more generally in the NE Pacific (Booth et al., 2008). Stomach contents of dissected specimens also indicate that *O. glabrum* is a deposit feeder, like members of the closely related genus *Ophiomusium* (Pearson and Gage, 1984). Single individuals generally occurred in the sediment around nodules, or under habitat-forming fauna such as non-stalked sponges and xenophyophores – giant, deep-sea foraminifera with delicate agglutinated tests (Goineau and Gooday, 2019).

The limited availability of food in the deep-sea soft-sediment benthos is a principal driver in structuring deep-sea benthic communities characterised by low population densities but high levels of diversity (Hardy et al., 2015). The CCZ is a vast, food-limited ecosystem, reliant on the arrival of particulate organic matter (POM) to the seafloor, dictated largely by productivity at the surface (Pennington et al., 2006). While nodule substrate availability plays a role in determining species distributions (Vanreusel et al., 2016), food availability – largely dependent on POM flux – will ultimately determine population densities. Having evolved under these relatively stable conditions, communities in the abyssal deep-sea may be poorly adapted and thus highly susceptible to future mining impacts. If, for example, elevated substrata provide a nursery-type habitat for *Ophiacantha cosmica*, then the ramifications for reproductive success following future mining disturbance events are cause for concern. In addition, filter-feeding organisms are expected to be severely impacted by plumes and settling aggregates generated by the resuspension of sea-floor sediments during nodule collections, resulting in long-term loss of suspension feeding organisms (Simon-Lledó et al., 2019). As with all deposit feeders, changes in surface-sediment chemical and physical characteristics following mining activity (e.g., remobilisation of formerly sequestered heavy metals and sediment compaction) could increase the toxicity of ingested sediments and impact *Ophiosphalma glabrum*’s ability to forage for food.

Food availability (i.e., POC flux) in soft-sediment abyssal habitats is subject to spatial variability both regionally, due to an undulating seafloor topography and at ocean-basin scales, due its dependency on surface productivity (Smith et al., 2008). Such heterogeneity in POC flux could create habitat networks of richer or poorer food supply over spatial scales at which reproductive kinetics and larval dispersal become ecologically relevant (Hardy et al., 2015). The CCZ lies along a POC flux gradient: lowest in regions that underlie central oligotrophic ocean gyres and highest beneath or adjacent to coastal and equatorial regions characterised by productive upwelling zones (Smith et al., 2008). Since population densities decrease concomitantly with POC flux (Tittensor et al., 2011), species in food impoverished areas often occur at densities below the minimum needed for reproductive success (e.g. critically low encounter rates for mating), rendering them wholly dependent on larval supply from populations in higher POC-flux habitats (the oligotrophic sink hypothesis, Hardy et al., 2015). The two species examined in the current study are relatively common in the CCZ and beyond (Booth et al., 2008; Amon et al., 2016; Christodoulou et al., 2019, 2020; Simon-Lledó et al., 2019). In fact, it is for this reason that the current reproductive study was even possible. However, the relative densities at which they occur at regional scales (such as between contract areas / APEIs) varies considerably (e.g. Christodoulou et al., 2020). Should mining activities proceed in regions of higher-POC flux that play a formative role as larval sources, this may have wide-reaching detrimental effects on the most food-limited populations of these and other species that employ planktonic larval dispersal.

Reproductive kinetics play a fundamental role in mediating biogeographic patterns, population dynamics, metapopulation connectivity and ultimately, species survival (Ramirez Llodra, 2002). While molecular approaches can be used to examine both historical and contemporary connectivity, the temporal resolution of these data remains too poor to discriminate between sporadic regional genetic exchange that then spreads locally over successive generations, and regular regional genetic exchange that acts to supplement local populations to a biologically meaningful degree. Reproductive ecology remains vital in bridging gaps in our understanding. The current study sought to add to the paucity of reproductive data for deep-sea species in nodule environments. It also highlights an uncomfortable truth in deep-sea ecology: that classical reproductive studies of this sort, which necessitate large numbers of individuals, may not be practicable in habitats such as the CCZ where the most susceptible species are typically the rarest. Support for more ambitious approaches that compensate for sampling constraints, such as temporal studies or permanent monitoring networks are urgently needed in order for conservation measures to be effective and informed.

## 6 Author Contributions

AH conceived reproductive study. AH, MC and PMA coordinated sample collection, fixation and preservation. MC and PMA performed all aspects of molecular ID. SRL performed specimen imagery, dissections, sectioning, histological, graphical and statistical analyses, prepared figures and wrote manuscript with scientific input from remaining authors.

## 8 Acknowledgments

We would like to acknowledge the captain and crew of the R/V Sonne and ROV Pilots for the Kiel 6000 (GEOMAR) and to the deep-sea scientists involved in cruise organisation as well as collection and processing of samples on board. Our thanks to António and Sandra Calado for the use of an ultramicrotome. A pre-print of this manuscript is available on bioRxiv, Laming et al. (2020) at https://doi.org/10.1101/2021.02.06.428832.

**9 References**

**9.1 Peer-reviewed**

**9.2 R software and package references (those not subject to peer review)**

## 10 Supplementary Material

The Supplementary Material for this article can be found in an accompanying PDF presentation entitled “Supplementary material.pdf”.

## 11 Data Availability Statement

Oocyte feret diameter measurements used in this study can be made available upon request. Molecular data and sample metadata may be found here: http://dx.doi.org/10.5883/DS-CCZ4. This dataset contains accession codes (Genbank), BOLD IDs, disc diameters, assigned sex and gonad state, along with photos, trace files and collection data.

## Footnotes

1 https://www.R-project.org/

2 https://inkscape.org

