## Supplementary figure 1 for "Comparative reproductive biology of deep-sea ophiuroids inhabiting polymetallic-nodule fields in the Clarion-Clipperton Fracture Zone"

bioRxiv
THE PREPRINT SERVER FOR BIOLOGY

### *Supplementary Material*

bioRχiv
THE PREPRINT SERVER FOR BIOLOGY

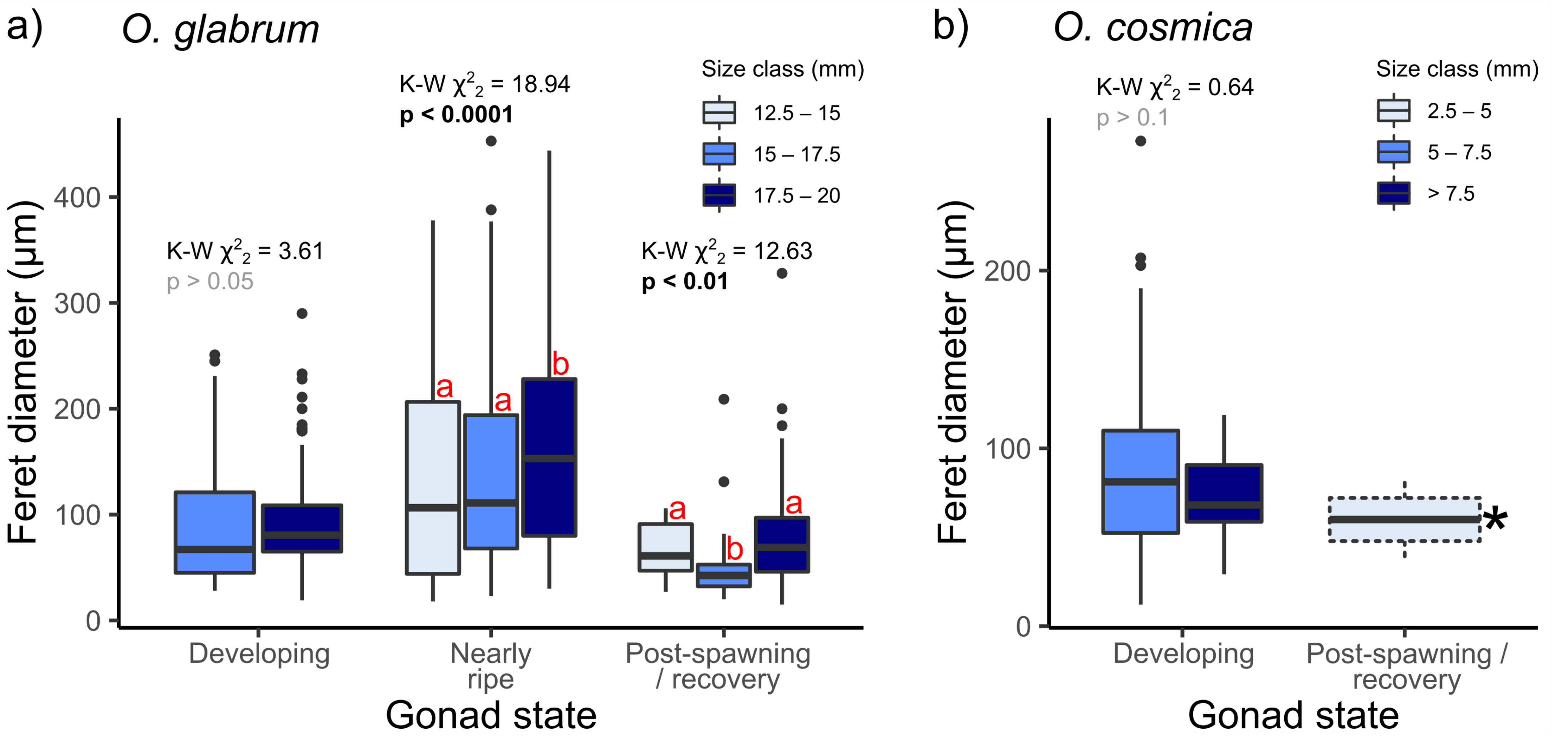


**Supplementary Figure 1. Overall size-class frequency distributions for both study species**
Analyses are based on twenty-five *O. glabrum* specimens, of which 14 were *nearly ripe*, 5 were *developing* and 6 were in *post-spawning / recovery* and seven *O. cosmica* specimens, of which all but one was *developing*. Box and whisker plots displaying variation in oocyte feret diameters as a function of binned disc-diameter data (respectively) for female specimens in each gonad state identified. Significant differences in oocyte feret diameter with size were assessed (where relevant) for each gonad state, using Kruskall-Wallis Chi-squared tests on ranked data. Significant post-hoc pairwise comparisons (Dunn's tests) are those with no letter annotations in common.
*Box and whisker values for this class based on two measurements only.
